# CAPHEINE, or everything and the kitchen sink: a workflow for automating selection analyses using HyPhy

**DOI:** 10.64898/2026.02.23.707482

**Authors:** Hannah Verdonk, Danielle Callan, Sergei L Kosakovsky Pond

## Abstract

**Summary:** Here we present CAPHEINE, a computational workflow that starts with a set of unaligned pathogen sequences and a reference genome and performs a comprehensive exploratory evolutionary analysis of the input data. CAPHEINE pairs nicely with studies of site-level selection dynamics, gene-level positive selection, and lineage-specific shifts in selective pressure. Our workflow is portable across Mac OS, Windows, and Linux, allowing researchers to focus on results.

**Availability and Implementation:** `CAPHEINE` is freely available at https://github.com/veg/CAPHEINE, along with a set of usage instructions.

## Introduction

Evolutionary dynamics and transmission mechanisms in viruses are incredibly diverse. This results in significant variability in their patterns of emergence, spread, and persistence (Pybus and Rambaut 2009; Leventhal et al. 2015). RNA viruses, such as Influenza A virus (IAV), Hepatitis C virus (HCV), Human Immunodeficiency Virus Type 1 (HIV-1), and SARS-CoV-2 are all characterized by rapid mutation rates and within-host evolution, allowing them to generate diverse viral variants within a host and generating strains with enhanced immune evasion or transmission potential (Luciani and Alizon 2009; Dingens et al. 2019; Lee et al. 2019). IAV, SARS-CoV-2, and HIV-1 are able to create further genetic diversity through reassortment or homologous recombination (Burke 1997; Jetzt et al. 2000; Steel and Lowen 2014).

Lack of genetic diversity can also play a role in viral evolution and dynamics. New infections are often characterized by transmission bottlenecks, as there is less diversity in the founder virus population than in the original host. The bottleneck can lead to genetic drift and the rise of potentially deleterious variants by chance alone (Markov et al. 2023). Additionally, vector-borne RNA viruses face stronger selective constraints due to their life cycle - dengue virus, for example, must persist in both vertebrate and invertebrate hosts (Holmes 2010; Nanaware et al. 2021).

Understanding how viruses spread and persist requires insight into their evolutionary dynamics. With vast amounts of viral sequence data available from sources such as GISAID and the Sequence Read Archive (SRA) (Shu and McCauley 2017; Katz et al. 2021), there is a pressing need for computational methods that can effectively probe this wealth of information and extract meaningful signals of selection. Such methods must be rapid, flexible, and scalable to accommodate frequent updates with newly available genomic data. A computational workflow is an ideal tool to analyze these data. Well-designed workflows must be documented, reproducible analyses; they are portable across different hardware platforms and operating systems, allowing researchers to focus on the high-level analysis logic and results without worrying about the specific software implementation, version control, or dependencies. Support for HPC schedulers and containerization platforms (*e*.*g*., Docker and Singularity) add flexibility - different analysis steps can be parallelized and isolated from the native OS environment.

Existing computational workflows to characterize viral evolution are often tailored to a particular viral pathogen or dataset, *e*.*g*., (Hadfield et al. 2018; Lucaci et al. 2022), and can be difficult to modify without technical experience. Other, more generalist workflows such as `V-pipe` and `ViralFlow v1.0` (Posada-Céspedes et al. 2021; da Silva et al. 2024) characterize positive selection on single nucleotide variants of interest within a single sample or population, but do not test for (and consequently, may not capture) selection across lineages or at conserved sites.

Here, we introduce the <u>C</u>omprehensive <u>A</u>utomated <u>P</u>ipeline using <u>H</u>yPhy for <u>E</u>volutionary <u>I</u>nference with <u>Ne</u>xtflow (CAPHEINE), a computational workflow that characterizes selection across sites, branches, and clades within a set of pathogen sequences. CAPHEINE is designed for exploratory hypothesis generation rather than definitive inference of adaptive mutations. Our workflow enables researchers to quickly profile sequence variation and establish correlations between selective pressure and amino acid composition. As a demonstration, we apply `CAPHEINE` to H5N1 IAV genomes isolated between 1975 and 2025 and identify signatures of selection across all viral genes. We also contrast selection across sites within the reservoir population of wild birds (Anseriformes and Charadriiformes) and within recent cattle spillover sequences from the 2025 outbreak, searching for signatures of viral adaptation to cattle hosts.

## Methods

We developed `CAPHEINE` with the goal of quickly and easily surveying the selection profile for a set of viral genes. Characterizing selection across the coding genome is a central analysis in evolutionary biology, and frequently occurs in studies that characterize genomic evolutionary dynamics, profile the strength and direction (positive or negative) of natural selection, connect evidence of coding sequence selective constraints with protein structure and function, and explore how shifts in selection pressure correlate with host changes (Langedijk et al. 2024; Siozios et al. 2024; Balech et al. 2025; Kotwa et al. 2025). In a recent paper in *PLOS Pathogens*, Huang *et al* characterize the antiviral mechanism of Zinc finger antiviral protein (ZAP) (Huang et al. 2024). The authors identify candidate viral interaction sites in mammalian ZAP orthologs by identifying sites with statistical evidence of positive selection. Subsequent *in vitro* mutagenesis at each candidate site revealed one mutant (N658A) with enhanced antiviral activity over the wild-type ZAP protein, confirming the functional importance of one candidate.

`CAPHEINE` streamlines the evolutionary discovery process by providing a single, standardized workflow that performs selection analyses quickly, reproducibly, and at scale (Figure 1). Many of the studies cited here have independently invested substantial effort in constructing custom pipelines for similar analyses, creating separate workflows that might be difficult to reproduce or compare across studies. `CAPHEINE` ensures consistency and transparency throughout all analyses by basing itself on the `nf-core` framework, a standardized set of tools and guidelines for writing bioinformatics pipelines in the `Nextflow` workflow language (Di Tommaso et al. 2017; Ewels et al. 2020). The `CAPHEINE` workflow takes just two input files: a “query” dataset comprising a single FASTA file containing unaligned full or partial pathogen genomes, and a “reference” dataset of gene sequences for the pathogen in fasta format (*e*.*g*., downloaded genomic coding sequences from NCBI RefSeq assembly `GCF_000864105.1`). As a result, users can begin running CAPHEINE with little or no additional data preparation.

**Figure 1.**
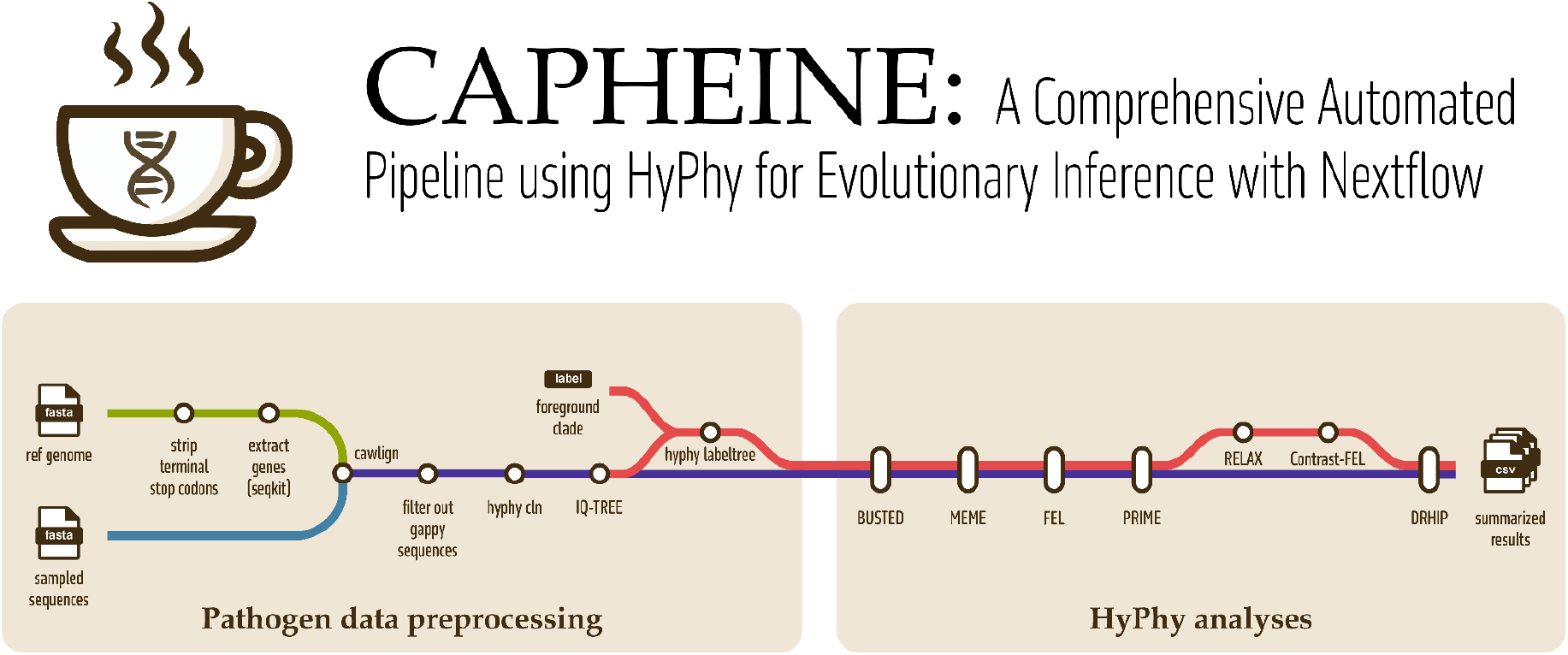
Overview of the `CAPHEINE` workflow, including data preprocessing steps and evolutionary analyses. Light green and blue lines represent two sources of input data: a reference genome and the pathogen sequences to be analyzed. Once the sequences are aligned to the reference genome (`cawlign`), the data undergoes further preprocessing and analysis. Optionally, a set of foreground taxa may also be provided, triggering the additional evolutionary analyses Contrast-FEL and RELAX (red line).

Optionally, users may also specify a subset of *foreground* sequences to test for differences in selection pressure against all other sequences. Contrasting the evolutionary history between two branch sets allows users to explore how selection intensity changes with lineage, as well as how individual sites evolve differently under different circumstances. The work of Kotwa *et al* provides a case study: the authors contrast selection in the S (spike) gene for a newly discovered *Eptesicus fuscus-*derived bat coronavirus and closely related sequences, and for a newly discovered *Myotis lucifugus-*derived bat coronavirus and closely related sequences (Kotwa et al. 2025). Using a combination of `HyPhy` methods, Kotwa *et. al* find that selection acts in a broadly similar manner for both the newly discovered coronaviruses and their most closely related sequences; there is no evidence for relaxed or intensified selection, and only one site in the *Eptesicus fuscus-*derived bat clade showed statistically significant evidence differential selection between the newly discovered coronavirus and its relatives.

To perform a similar analysis, taxa (or branches) can be marked as foreground lineages with either a list of newline-separated sequence identifiers, or with a regular expression pattern that can be used to match the foreground sequence identifiers. `CAPHEINE` will report any relaxation or intensification of selection between foreground and reference lineages, as well as site-specific differences in selection pressure and amino acid composition between foreground and reference. The foreground taxa need not be monophyletic or consistently present across all genes.

`CAPHEINE` first trims the terminal stop codons (if needed) from the reference sequences and removes query sequences with more than 50% gaps or ambiguous bases (Figure 1). Then, query sequences are mapped to each reference gene using `cawlign v0.1.14 (-s BLOSUM62 -t codon`). Cawlign is a codon-aware pairwise alignment algorithm based on dynamic programming with frame-preserving gap penalties and explicit modeling of codon triplets, ensuring reading frame integrity during alignment (Anon 2025). Duplicate sequences are removed from each gene’s alignment using `hyphy cln` , which speeds up computation without affecting any `hyphy` methods’ estimate of selection. A maximum likelihood tree is then inferred for each alignment using `IQ-Tree2 v2.4.0 (-T 8 -m GTR+I+G`) (Minh et al. 2020). Although model selection (*e*.*g*., ModelFinder) could be performed, we prioritize rapid tree estimation for large datasets using the GTR+I+G model, as HyPhy subsequently re-estimates branch lengths under codon models appropriate for each downstream analysis. If a foreground clade is provided, internal branches of the foreground clade are labeled as “Foreground” and internal branches of the background clade are labeled as “Reference” using the `label-tree` function in `HyPhy v2.5.84` (`--internal-nodes ‘All descendants’ --leaf-nodes ‘Skip’`) (Kosakovsky Pond et al. 2020). Optionally, users can label all branches (including leaf branches) belonging to the foreground and reference clades, respectively.

We have selected six molecular evolutionary analyses (Spielman et al. 2019), implemented in the HyPhy package, as the core of our workflow (Table 1). These analyses were chosen because they are popular and frequently used in other studies (West et al. 2021; Huang et al. 2024; Policarpo et al. 2024; Balech et al. 2025; Drabeck et al. 2025). Note that Contrast-FEL and RELAX are only run if a foreground set of branches is provided. Because site-level methods test hundreds to thousands of codons per gene, users are strongly encouraged to apply multiple testing correction (*e*.*g*., Benjamini–Hochberg FDR) when interpreting results.

**Table 1.** A summary of all the evolutionary methods included in CAPHEINE. Contrast-FEL and RELAX are only run if a foreground clade is provided.

| Method | Tests for | Details | Citation |
| --- | --- | --- | --- |
| FEL | pervasive (*i.e.*, phylogeny-wide) diversifying selection at each site in an alignment. | FEL is used as a discovery tool to identify any site that may be under selection. Multiple testing correction is not applied to p-values. | (Kosakovsky Pond and Frost 2005) |
| MEME | episodic diversifying selection - *i.e.*, diversifying selection that has occurred on only a single branch, or subset of branches in the phylogeny along a site. | MEME is used as a discovery tool to identify any site that may be under selection. Multiple testing correction is not applied to p-values. | (Murrell et al. 2012) |
| BUSTED | gene-wide evidence of positive episodic diversifying selection | Statistical significance is assessed by LRT and reported at the gene level. | (Murrell et al. 2015) |
| PRIME | whether changes in amino acid properties (e.g., hydrophobicity) at a site are favored or disfavored | We have chosen to focus on the set of five biochemical properties described by Atchley *et. al.* (Atchley et al. 2005) when assessing property conservation at each site. | Forthcoming |
| Contrast-FEL | Whether a site evolves differently on a specified foreground clade, relative to the set of reference branches | Multiple testing correction is not applied to p-values. Some sites are likely false positives. | (Kosakovsky Pond et al. 2021) |
| RELAX | whether the strength of selection has been relaxed or intensified gene-wide on a specified foreground clade, relative to the set of reference branches | RELAX is **not** designed to test a set of lineages for the presence or absence of diversifying selection, but to find relaxation ($K < 1$) or intensification ($K > 1$) of selection on the foreground branches relative to the reference branches. | (Wertheim et al. 2015) |

### Case study: H5N1 host shift

Here, we demonstrate how to run CAPHEINE and interpret the output using H5N1 IAV viral sequences collected between 1975 and August 12, 2025 and uploaded to GenBank. Running CAPHEINE with default parameters allows us to explore broad, gene- and site-level patterns of selection across our entire dataset, so that we can see if the patterns we find are consistent with what we already know about IAV evolutionary dynamics. However, we also wish to find sites that might be under differential selection due to the host shift from the wild bird reservoir population to the cattle outbreak population. To that end, we specify cattle isolates as our foreground taxa with the additional argument `--foreground_regexp “cattle”`; the regular expression will match all fasta IDs that contain the string “cattle” and mark those sequences as foreground.

Our starting dataset included 28,751 genomic sequences from over 2,706 cattle isolates and 41,988 genomic sequences from over 3,396 wild bird (*Anseriformes* and *Charadriiformes*) isolates. We are unsure of the exact number of isolates due to isolate naming inconsistencies in Genbank, so we report the number of isolates with exactly 8 associated segments per dataset. NCBI’s RefSeq datasets provide users with the option of downloading reference coding sequences for annotated genomes; we took advantage of this feature to obtain H5N1 IAV reference gene sequences (NCBI RefSeq assembly `GCF_000864105.1`). We excluded the short overlapping PA-X and PB1-F2 genes from our analyses. After duplicate removal with `hyphy cln`, each gene alignment contained between 980 and 4600 unique sequences (Table S1).

We restricted the `HyPhy` analyses to include only *internal branches* to capture fixed differences due to selection for transmission and survival, rather than intra-host evolution. Users may configure `CAPHEINE` to analyze all branches if desired. Analyses also assume the Universal genetic code for all aligned sequences, but this can be easily reconfigured.

`CAPHEINE` summarizes and reports all analysis results as plain-text CSV files. These files can be readily opened in common spreadsheet programs, allowing users to easily inspect or explore the data without requiring specialized software or advanced computational expertise. Results files also drop directly into downstream analytics in `R, Python`, or visualization notebooks, enabling rapid iteration from first pass results to figures suitable for reports and manuscripts.

Our first question was “*How are genes and sites evolving across the entire dataset, and is the evolutionary pattern we find consistent with what we already know about IAV*?” We confirm a few expected evolutionary patterns: most genes showed evidence of strong purifying selection, with mean ⍵ values in the range 0.0627 - 0.8193 for the foreground (cattle) internal branches and many individual sites evolving under negative selection (Figure 2, Table S2). However, when we filter for genes with significant `BUSTED` results, we find that over half of all gene products (PB2, PB1, PA, NA, NS1) also showed some statistical evidence of episodic diversifying selection for at least one site on at least one branch. At the site level, there is evidence that one or more biochemical properties from the set containing secondary structure, bipolar, molecular volume, amino acid composition, and electrostatic charge are under selection for at least one site in each gene.

**Figure 2.**
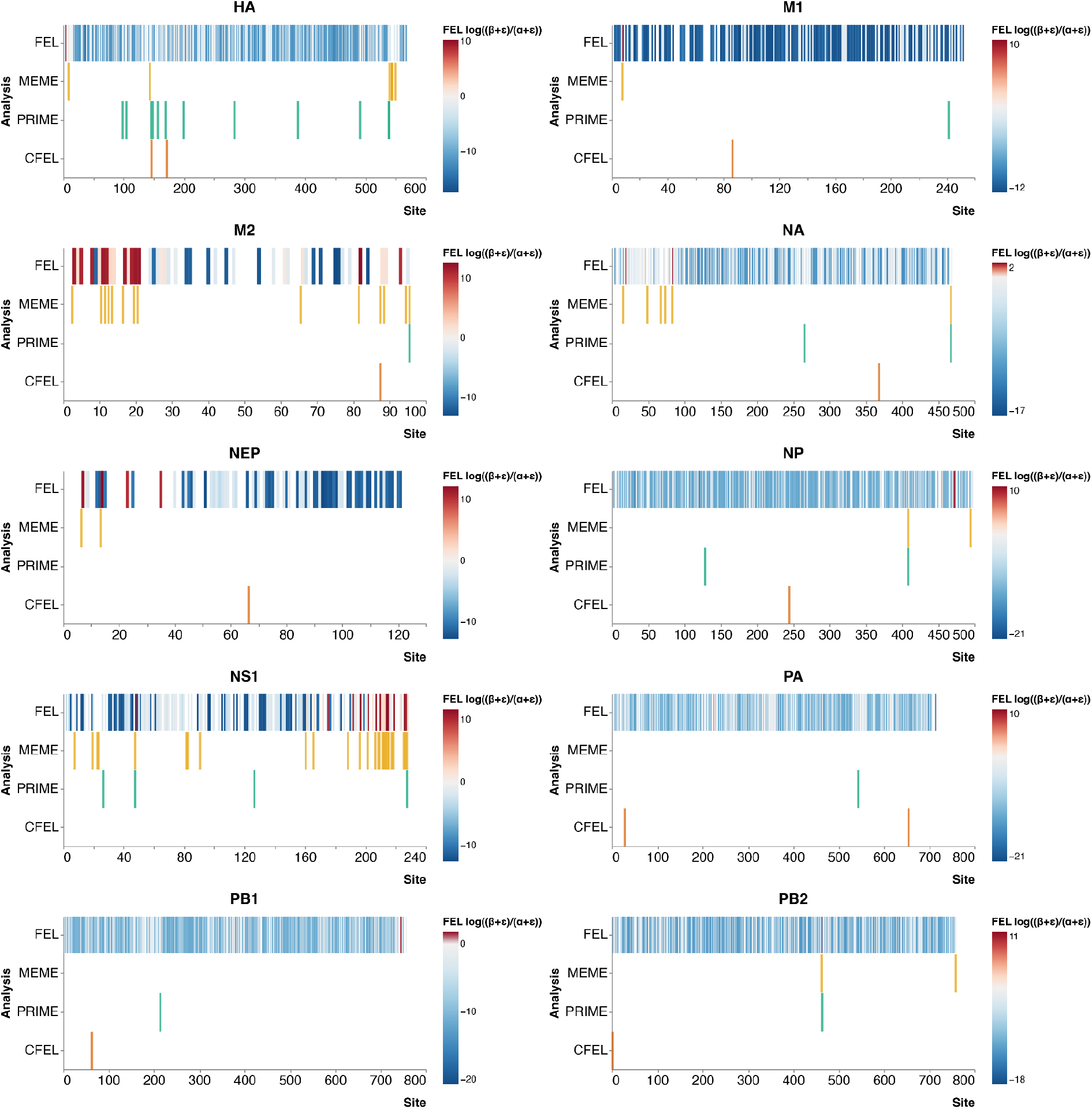
Site-wise CAPHEINE results. Significant sites for each method are shown as colored bars,while insignificant sites are left blank. For FEL, direction of selection (positive or negative) is shown as a heatmap color value, with red indicating positive selection and blue indicating negative selection for significant sites.

Our second question was “*How does IAV evolution differ between the wild bird reservoir hosts and the cattle outbreak*?” RELAX analysis suggests that H5N1 sequences in cattle hosts are under **intensified** selection in HA, NS1, and PB2 and under **relaxed** selection in NP, PA, and PB1, relative to the viral sequences found in wild bird hosts (Figure 3). Our findings have some precedent: Misra *et. al* independently identified relaxed selection in PB2 internal branches in the entire 2.3.4.4b clade of highly pathogenic avian influenza, relative to other H5N1 clades (Misra et al. 2024). We also filtered our results for individual sites with evidence of intensified positive selection; *i*.*e*., sites with significant MEME test results for episodic positive selection and significant Contrast-FEL test results with β_cattle_ > β_wild birds_. We identified two sites meeting both criteria in PA and M2 (Table S3, starred sites). Of these, only site 88 of the M2 gene had evidence of both intensified positive selection and a different majority residue among the cattle branches (asparagine) and the wild bird branches (aspartate), possibly indicating a functional change in the protein product. The M2 gene encodes matrix protein 2, a component of the viral envelope involved in viral genome packaging and the production of infectious viral particles (Hughey et al. 1995; Chen et al. 2008; Ozawa et al. 2009). M2 residues 82 to 89 are specifically required to produce larger titers of infectious virus (McCown and Pekosz 2006), and residues in this region were shown to be under positive selection in human influenza as well (Furuse et al. 2009).

**Figure 3.**
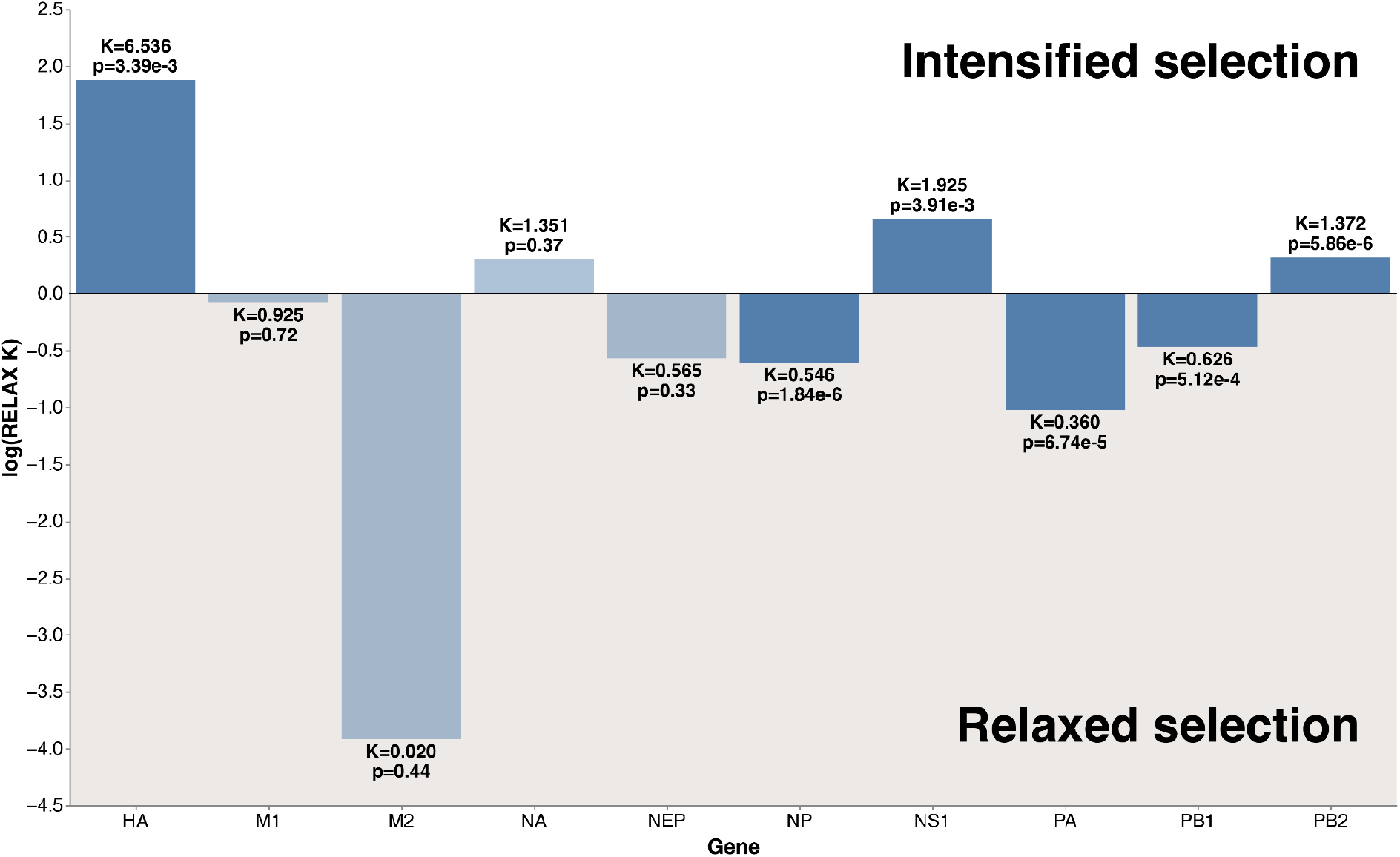
RELAX results for H5N1 sequences. The K-value is the Foreground relaxation/intensification parameter from the RELAX analysis, plotted here on a log scale. The K-value for the Foreground branches is estimated relative to the Reference branches, where K is set to 1 (not shown). Lower K-values (log(K) < 0) indicate relaxed selection compared to the Reference branches, while higher K-values (log(K) > 0) indicate intensified - but not *necessarily* positive - selection. p < 0.05 indicates that the differences are significant (dark blue bars).

### Usage and implementation

After installing Nextflow, `CAPHEINE` may be run with default parameters using the following command:

~~~
      nextflow run veg/CAPHEINE \
          -profile <docker/singularity/conda/custom> \
          --reference_genes <reference_genes.fasta> \
          --unaligned_seqs <unaligned_seqs.fasta> \
          --outdir <OUTDIR>
~~~

CAPHEINE requires two primary inputs: (i) a FASTA file containing unaligned pathogen genomes and (ii) a FASTA file of reference coding sequences. Optional arguments allow users to specify foreground lineages via a newline-separated list or regular expression pattern. More detailed usage instructions can be found at https://github.com/veg/CAPHEINE. Output consists of per-gene and per-site CSV files summarizing likelihood ratio statistics, parameter estimates, and p-values for each HyPhy method. Output files can be viewed in any spreadsheet software package or visualized at https://observablehq.com/@hverdonk/drhip-result-viewer

## Discussion

`CAPHEINE` streamlines exploratory selection evolutionary analysis by bundling the most commonly used `HyPhy` tests for selection into a single, coherent workflow. With `CAPHEINE`, researchers can ask — and rigorously answer — routine questions about gene- and site-specific pathogen evolution that would otherwise require bespoke scripting and repeated data wrangling. The integrated analysis suite also supports comparative questions across host reservoirs or epidemic waves, identifying relaxed or intensified selection across genes and pinpointing specific sites with altered biochemical constraints. In our H5N1 case study, `CAPHEINE` identified statistically significant evidence of episodic selection in the HA viral surface protein and in multiple RNA polymerase subunits. Our workflow easily identified specific sites with host-associated shifts in selection pressure, one of which (site 88 in gene M2) had evidence of intensified positive selection in cattle-associated viruses and an associated change in the majority amino acid residue. Such a change could indicate how the virus is adapting to thrive in a new host.

A key strength of `CAPHEINE` is practical usability. Because the pipeline is distributed with Conda, `Docker`, and `Singularity` options, it runs reproducibly on laptops, HPC clusters, and containerized cloud environments without manual environment tuning. `CAPHEINE` benefits from the standard configuration patterns, clear provenance, and community-tested orchestration of Nextflow and nf-core, and the workflow’s modular architecture enables robust process management and scalable parallelism. From the user’s perspective, inputs are minimal: just the sequences to analyze and a set of reference coding genes (which can frequently be found within the NCBI genome datasets), plus an optional foreground list or regular expression for contrastive selection tests.

`CAPHEINE` is agnostic to pathogenicity and can be applied to non-viral, non-pathogenic microbes so long as coding sequences and suitable references are provided. However, users should be aware that CAPHEINE does not account for paralogs among the reference genes, non-haploid genomes, or species-specific idiosyncrasies — the handling of these is left to the discretion of the species expert. We recommend exploring population genomic methods for evolutionary testing of non-haploid species. In addition, CAPHEINE does not currently perform automated recombination screening (*e*.*g*., via GARD), as such analyses can become computationally prohibitive for very large datasets. For segmented viruses such as Influenza A virus, reassortment is naturally accommodated by analyzing each segment independently, and homologous recombination within segments is rare. However, for recombining pathogens such as HIV-1 or coronaviruses, users should perform recombination screening prior to running selection analyses, as recombination can inflate false positive rates in site-level tests (Kosakovsky Pond et al. 2006).

`CAPHEINE` standardizes pathogen sequence preprocessing and analysis via a comprehensive selection toolkit. Researchers and surveillance teams can analyze new isolates quickly to gain insight about selective pressures, functional hotspots, and lineage-specific shifts. `CAPHEINE` grants researchers a window into viral evolutionary dynamics, providing actionable leads for experimental follow-up, vaccine and drug target prioritization, and public health decision-making.

## Acknowledgements

We are grateful to the nf-core community for developing and maintaining the tools used to build CAPHEINE.

