## supplemental tables for "CAPHEINE, or everything and the kitchen sink: a workflow for automating selection analyses using HyPhy"

### Supplement

| **Table S1.** Number of H5N1 sequences per gene after removing duplicates and sequences with more than 50% gaps or ambiguous nucleotides. | |
|---|---|
| **Gene** | **Sequences** |
| PB2 | 4311 |
| PB1 | 4190 |
| PA | 4284 |
| NP | 3390 |
| NA | 4047 |
| HA | 4594 |
| M2 | 980 |
| M1 | 2076 |
| NEP | 1430 |
| NS1 | 2574 |

**Table S2.** Overall selection characterization of H5N1 internal branches. **sites**: the number of codon sites in the alignment. Sites under positive selection have been inferred using MEME, negative selection using FEL, clade-specific $\omega_{clade} > \omega_{other\ clade}$ using Contrast-FEL. **BUSTED EDS**: the p-value of a test for episodic diversifying selection (EDS) on internal branches and the percent of branch sites for which $\omega \geq 1$. **branches**: the host-specific number of non-zero length branches included for selection testing. **Tree length**: the cumulative length of all branches assigned to each host (contrast-FEL, scaled in expected substitutions/nucleotide site). **ω**: mean estimate on host-specific internal branches (MG94xREV model).

| Gene | sites | Sites evolving under (p≤0.05) | | | BUSTED EDS (% of branch sites) | Host | branches | tree length | ω |
|---|---|---|---|---|---|---|---|---|---|
| | | positive | negative | clade-specific | | | | | |
| **HA** | 568 | 22 | 466 | 2 | N.S. | **Cattle (foreground)** | 139 | 0.1012 | 0.2649 |
| | | | | | | **Wild birds (reference)** | 1799 | 3.7689 | 0.1356 |
| **M1** | 252 | 1 | 192 | 1 | N.S. | **Cattle (foreground)** | 46 | 0.0612 | 0.0625 |
| | | | | | | **Wild birds (reference)** | 635 | 1.7726 | 0.0533 |
| **M2** | 97 | 19 | 20 | 1 | N.S. | **Cattle (foreground)** | 7 | 0.0138 | 1.1916 |
| | | | | | | **Wild birds (reference)** | 267 | 1.1929 | 0.7929 |
| **NA** | 469 | 19 | 363 | 1 | 0.0011561 (2.18%) | **Cattle (foreground)** | 123 | 0.1141 | 0.2344 |
| | | | | | | **Wild birds (reference)** | 1462 | 4.2578 | 0.1862 |
| **NEP** | 121 | 5 | 58 | 1 | N.S. | **Cattle (foreground)** | 19 | 0.0569 | 0.3941 |
| | | | | | | **Wild birds (reference)** | 397 | 2.0617 | 0.3061 |
| **NP** | 498 | 6 | 428 | 1 | N.S. | **Cattle (foreground)** | 102 | 0.0801 | 0.1506 |
| | | | | | | **Wild birds (reference)** | 1210 | 2.6860 | 0.0444 |
| **NS1** | 230 | 33 | 116 | 0 | 0.0001892 (12.63%) | **Cattle (foreground)** | 71 | 0.1098 | 0.7334 |
| | | | | | | **Wild birds (reference)** | 800 | 3.0148 | 0.3172 |
| **PA** | 716 | 10 | 599 | 2 | 7.22E -16 (1.20%) | **Cattle (foreground)** | 166 | 0.1012 | 0.2048 |
| | | | | | | **Wild birds (reference)** | 1591 | 2.9522 | 0.0912 |

| | | | | | | | | | |
|---|---|---|---|---|---|---|---|---|---|
| **PB1** | 757 | 6 | 662 | 1 | 0.0002864 (0.85%) | **Cattle (foreground)** | 181 | 0.0944 | 0.1390 |
| | | | | | | **Wild birds (reference)** | 1582 | 2.9559 | 0.0631 |
| **PB2** | 759 | 16 | 673 | 1 | 0 (0.82%) | **Cattle (foreground)** | 196 | 0.1055 | 0.1837 |
| | | | | | | **Wild birds (reference)** | 1605 | 3.6323 | 0.0600 |

**Table S3.** Individual sites which show positive selection on the H5N1 cattle host internal branches (MEME p-value ≤ 0.05), or where selection is operating differently on H5N1 cattle host internal branches compared to H5N1 wild bird host internal branches (Contrast-FEL p-value ≤ 0.05). **Codon**: codon position in the multiple sequence alignment; **pos**: MEME p-value; **dif.**: Contrast-FEL p-value; **composition**: amino-acid composition of cattle and wild bird sequences at this site; **substitutions**: inferred substitutions on cattle and wild bird clade internal branches at this site. *: a site is marked with * if it is **both** positively selected and intensified ($\beta_{cattle} > \beta_{wild\ birds}$) in cattle sequences; #: the majority residue is different between cattle and wild bird sequences. '-' : test result is not significant, or no substitutions occurred within a group

| Gene | Codon | pos. | dif. | PRIME | Composition | | Substitutions | |
|---|---|---|---|---|---|---|---|---|
| | | | | | Cattle | Wild birds | Cattle | Wild birds |
| **HA** | 10 | 0 | - | - | I/39 -/5 M/2 ?/1 | I/1992 M/30 V/14 -/136 T/440 L/2 A/232 ?/2 K/1 S/3 | I:M/1 | I:M/5 I:T/4 I:V/2 A:T/1 S:T/1 M:V/1 |
| **HA** | 14 | 0.016 | - | - | V/43 -/3 ?/1 | V/2712 -/116 A/13 ?/3 F/1 G/2 I/1 L/1 T/3 | - | A:V/2 V:V/17 G:V/1 T:V/1 |
| **HA** | 63 | 0.029 | - | - | V/46 ?/1 | V/2826 ?/13 I/10 K/1 H/1 L/1 | - | V:V/5 I:V/2 |
| **HA** | 87 | 0.02 | - | - | I/34 -/1 L/12 | I/2202 N/9 T/291 ?/2 L/337 F/3 -/6 V/1 D/1 | - | I:T/14 I:I/1 I:L/2 L:L/6 F:L/1 T:T/1 N:T/1 I:N/1 |
| **HA** | 99 | 0.047 | - | overall, composition | A/35 I/12 | A/2347 T/29 D/155 S/32 P/2 I/276 N/1 V/2 -/6 ?/2 | - | A:S/1 A:D/7 D:D/3 A:A/5 A:I/1 I:I/1 A:T/5 |
| **HA** | 136 | 0.027 | - | - | S/37 N/10 | S/2010 N/110 ?/10 I/3 G/4 T/1 D/708 E/6 | N:S/1 | S:S/4 N:S/6 G:S/1 D:N/7 D:S/1 D:D/4 D:E/1 |
| **HA** | 144 | 0.002 | - | overall, composition | S/35 -/12 | S/2694 -/139 N/1 T/14 ?/1 F/2 P/1 | - | S:T/2 S:S/1 F:S/1 |

| | | | | | | | | |
|---|---|---|---|---|---|---|---|---|
| **HA** | 146 | 0.005 | - | overall | G/47 | G/2834 -/7 P/9 R/1 W/1 | - | G:G/13 G:R/1 P:R/1 |
| **HA** | 147 | - | 0.001 | - | V/44 M/3 | V/2840 M/3 E/1 -/7 A/1 | - | V:V/4 |
| **HA** | 156 | 0.004 | - | - | A/35 R/12 | A/1475 V/43 S/7 T/105 K/161 Q/30 R/355 N/648 ?/1 M/13 -/7 G/2 I/4 H/1 | - | A:V/5 A:A/6 A:T/3 K:T/2 K:Q/1 K:M/1 K:K/1 K:R/5 R:R/4 I:R/1 K:S/1 N:S/2 K:N/1 N:N/1 M:T/2 T:V/1 T:T/1 A:M/1 |
| **HA** | 172 | - | 0.018 | overall, volume | A/41 T/6 | A/2421 E/1 V/3 T/400 -/5 S/18 ?/1 P/1 G/1 I/1 | A:T/2 | A:A/7 A:V/1 A:T/7 A:S/2 T:T/1 |
| **HA** | 178 | 0.036 | - | - | I/33 V/2 K/3 R/9 | I/1514 V/2 R/517 M/10 K/794 S/8 -/5 G/1 T/1 | I:V/1 | I:M/3 I:R/2 R:R/2 R:S/2 K:R/5 |
| **HA** | 204 | 0.033 | - | - | T/47 | T/2740 I/86 A/8 K/3 -/6 E/1 M/5 V/3 | - | I:T/12 T:T/3 I:V/1 I:M/1 K:T/1 |
| **HA** | 205 | 0.004 | - | overall, bipolar, structure | N/35 R/12 | N/1490 K/513 S/1 R/810 Q/13 -/6 A/1 M/5 T/4 G/3 D/3 E/2 ?/1 | - | K:N/7 K:R/17 K:K/6 K:Q/3 R:R/1 R:T/1 G:R/1 M:R/1 |
| **HA** | 344 | 0.009 | - | overall, bipolar | K/47 | K/2717 -/89 G/1 A/2 R/42 I/1 | - | E:K/2 E:G/1 A:E/1 K:K/1 K:R/4 |
| **HA** | 413 | 0.038 | - | volume, composition, charge | G/47 | G/2825 ?/23 R/1 E/3 | - | G:G/8 E:G/1 |
| **HA** | 418 | 0.012 | - | - | N/47 | N/2710 ?/31 T/3 K/4 S/99 A/2 R/2 C/1 | N:N/1 | N:N/8 A:N/1 N:S/5 |
| **HA** | 540 | 0 | - | overall, bipolar, volume, composition, charge | A/43 -/3 T/1 | A/2787 -/46 S/1 C/10 V/3 T/5 | - | A:A/6 A:C/1 A:T/1 |
| **HA** | 541 | 0.004 | - | overall | S/44 -/3 | S/2795 -/46 E/10 T/1 | - | S:S/3 E:S/1 |
| **HA** | 544 | 0 | - | composition | A/42 -/4 V/1 | A/2125 -/52 T/11 S/11 V/648 E/1 ?/2 M/1 G/1 | - | A:T/3 A:S/1 A:V/4 V:V/2 A:A/1 |
| **HA** | 545 | 0 | - | overall, structure, volume, composition, charge | L/43 -/4 | L/2784 ?/1 -/54 G/1 T/9 S/1 R/1 P/1 | - | L:L/3 L:T/1 |
| **HA** | 546 | 0.003 | - | - | A/43 -/4 | A/2778 -/55 ?/5 V/1 T/1 G/10 E/2 | - | A:G/1 |

| | | | | | | | | |
|---|---|---|---|---|---|---|---|---|
| **HA** | 550 | 0 | - | overall | A/41 -/6 | A/2764 T/2 -/72 L/1 S/12 D/1 | - | A:S/1 |
| **HA** | 552 | 0.031 | - | composition | L/41 -/6 | L/2772 -/73 ?/1 M/3 I/2 H/1 | - | L:L/5 L:M/1 |
| **M1** | 8 | 0 | - | overall | E/103 -/3 | E/1249 -/15 R/3 ?/1 | - | E:R/1 |
| **M1 #** | 87 | - | 0.051 | overall, composition | N/20 T/86 | N/1190 S/1 T/76 A/1 | N:T/2 T:T/1 | N:N/3 |
| **M2** | 3 | 0.008 | - | volume | I/9 ?/3 | I/714 T/20 | - | I:T/2 |
| **M2 #** | 4 | 0.05 | - | - | F/1 S/7 ?/4 | S/290 F/441 L/3 | - | F:S/2 S:S/1 F:L/1 |
| **M2** | 8 | 0.042 | - | - | C/12 | C/722 Y/11 G/1 | - | C:Y/2 |
| **M2** | 10 | 0.038 | - | - | P/12 | P/604 H/107 L/22 S/1 | - | H:P/3 L:P/3 |
| **M2** | 11 | 0.006 | - | - | T/12 | T/719 I/14 S/1 | - | I:T/3 |
| **M2 #** | 12 | 0.016 | - | - | K/12 | K/284 R/450 | - | K:R/3 |
| **M2** | 13 | 0.001 | - | overall | N/12 | N/677 T/25 K/22 D/2 S/6 H/2 | - | N:T/4 K:N/3 H:N/1 N:S/1 |
| **M2 #** | 14 | 0.005 | - | volume | G/12 | G/347 E/387 | - | E:G/7 |
| **M2** | 17 | 0.019 | - | - | C/12 | C/717 Y/15 G/1 ?/1 | - | C:Y/4 |
| **M2** | 19 | 0.042 | - | - | C/12 | C/702 S/1 Y/31 | - | C:Y/3 |
| **M2** | 20 | 0.005 | - | - | S/12 | S/706 R/4 N/19 I/4 G/1 | - | R:S/1 N:S/4 I:S/1 |
| **M2** | 21 | 0.013 | - | - | D/12 | D/692 G/42 | - | D:G/5 |
| **M2 #** | 28 | 0.033 | - | - | I/12 | I/337 ?/1 T/4 V/379 F/7 S/1 A/3 L/2 | - | I:T/1 I:V/5 F:V/1 I:L/1 |
| **M2** | 66 | 0.028 | - | - | E/12 | E/633 G/13 -/2 K/6 A/78 T/2 | - | E:G/4 E:E/1 E:K/2 A:E/1 |
| **M2** | 82 | 0 | - | - | S/12 | S/561 N/163 C/3 I/1 -/3 ?/1 D/2 | - | N:S/9 C:S/1 D:S/1 |
| **M2 *#** | 88 | 0.018 | 0.177 | - | N/11 -/1 | D/662 N/26 -/25 Y/7 ?/1 V/1 G/9 K/1 E/1 M/1 | D:N/2 | D:N/4 D:G/1 D:Y/2 |

| | | | | | | | | |
|---|---|---|---|---|---|---|---|---|
| **M2** | 89 | 0.008 | - | - | G/8 -/3 C/1 | G/643 -/27 S/43 D/13 V/6 C/1 A/1 | C:G/1 | G:G/2 G:V/1 G:S/6 D:G/3 C:G/1 |
| **M2** | 95 | 0.026 | - | bipolar, composition | E/8 -/4 | E/665 -/61 P/1 V/1 K/2 G/2 Q/1 N/1 | - | E:E/9 E:N/1 |
| **M2** | 96 | 0 | - | overall, bipolar, structure, volume, composition, charge | L/8 -/4 | L/664 -/68 K/1 E/1 | - | K:L/1 L:L/5 E:L/1 |
| **NA** | 2 | 0.006 | - | - | N/128 -/4 | N/2410 -/65 E/6 D/2 Y/5 M/1 V/1 H/1 S/15 ?/15 | - | E:N/1 N:S/1 |
| **NA** | 16 | 0.002 | - | overall, volume, charge | V/126 I/1 -/1 A/2 G/2 | V/2287 A/82 ?/2 -/26 I/104 M/1 T/17 G/2 | G:V/1 | A:V/15 V:V/6 I:V/7 I:T/1 G:V/1 |
| **NA** | 19 | 0.019 | - | - | I/130 -/1 M/1 | I/2362 T/44 -/25 L/1 V/41 M/48 | - | I:T/6 I:I/2 I:M/6 I:V/2 |
| **NA** | 42 | 0.017 | - | overall, volume, composition | N/112 V/1 F/19 | N/2232 K/2 T/2 S/29 -/17 D/7 H/2 V/35 F/178 L/1 A/15 I/1 | F:N/1 N:N/1 | N:N/5 S:S/2 N:S/2 D:N/1 H:N/1 N:V/1 V:V/1 F:V/2 A:V/2 |
| **NA** | 46 | 0.045 | - | - | P/112 T/20 | P/1598 L/3 S/76 H/4 -/22 A/414 V/63 G/3 T/286 ?/1 D/47 N/3 E/1 | P:T/1 | P:S/7 H:P/1 A:P/1 A:V/3 A:T/5 A:A/1 A:D/2 D:N/1 D:G/1 A:S/1 P:P/1 |
| **NA** | 49 | 0 | - | overall, bipolar | C/112 -/20 | C/1681 -/836 N/1 T/1 F/1 Y/1 | C:S/1 | C:S/1 N:S/1 N:T/1 S:T/1 C:C/1 |
| **NA** | 57 | 0.038 | - | - | E/112 -/20 | E/1673 G/6 -/838 K/4 | - | D:E/1 E:E/3 E:K/1 E:G/1 |
| **NA** | 60 | 0.032 | - | - | T/112 -/20 | T/1680 I/1 -/838 G/2 | - | T:T/5 G:T/1 |
| **NA** | 68 | 0 | - | - | N/110 -/20 E/1 D/1 | N/1554 S/195 H/3 K/2 -/764 R/2 T/1 | E:S/1 | N:S/2 H:N/1 Q:R/1 N:T/1 S:T/1 |
| **NA** | 72 | 0.004 | - | - | T/130 S/1 I/1 | T/2469 N/5 I/8 -/10 E/2 A/27 | - | T:T/7 E:T/1 N:T/1 A:T/1 |
| **NA** | 74 | 0.002 | - | - | L/8 F/124 | F/2237 -/10 L/153 V/4 Y/2 S/19 C/60 P/36 | - | V:V/1 F:S/3 F:F/2 C:F/2 F:P/2 F:L/1 |
| **NA** | 77 | 0.035 | - | overall, bipolar, structure | E/130 K/1 D/1 | E/2425 A/1 ?/4 G/35 K/11 D/35 -/9 R/1 | - | E:G/4 E:E/11 E:K/2 E:Q/1 D:E/2 |
| **NA** | 84 | 0 | - | - | A/19 T/113 | T/2224 I/17 R/1 A/160 Y/2 K/106 -/7 M/1 V/1 P/2 | A:T/1 | I:T/4 A:T/6 K:T/1 T:T/2 P:T/1 |

| | | | | | | | | |
|---|---|---|---|---|---|---|---|---|
| **NA** | 258 | 0.047 | - | - | I/110 M/21 V/1 | I/1811 M/689 V/18 A/2 L/1 | - | I:M/11 I:V/2 M:V/1 |
| **NA** | 263 | 0.039 | - | - | V/132 | V/2510 I/10 A/1 | V:V/1 | V:V/9 |
| **NA** | 366 | 0.022 | - | - | S/111 -/2 N/19 | S/1653 N/674 I/6 R/1 T/2 V/2 H/165 -/4 D/3 G/10 ?/1 | - | N:S/4 N:N/3 N:V/1 H:N/1 D:N/1 G:S/2 S:S/1 |
| **NA** | 369 | - | 0.031 | overall | S/116 -/2 I/4 V/1 N/5 R/4 | S/2439 I/25 N/40 R/12 -/4 G/1 | I:S/1 N:S/1 R:S/1 | I:S/2 S:S/11 N:S/2 R:S/3 |
| **NA** | 435 | 0.032 | - | - | T/130 -/2 | T/2502 -/17 A/2 | T:T/1 | T:T/8 |
| **NA** | 468 | 0 | - | overall | D/128 -/3 N/1 | D/2398 -/106 T/7 ?/7 G/1 N/1 A/1 | - | D:T/1 |
| **NA** | 469 | 0.009 | - | - | K/128 -/4 | K/2382 N/8 -/116 R/2 S/6 Q/1 ?/5 E/1 | K:K/1 | K:N/1 K:S/1 K:K/1 |
| **NEP** | 7 | 0.002 | - | overall | - | L/1006 ?/1 P/58 -/1 I/3 T/1 F/2 H/1 | - | L:P/5 |
| **NEP** | 14 | 0 | - | - | - | M/727 V/239 L/2 T/5 I/6 A/40 G/29 E/1 K/2 Q/21 ?/1 | - | M:V/7 I:M/1 G:V/2 A:V/2 A:M/1 M:Q/1 |
| **NEP** | 60 | 0.035 | - | - | - | S/650 N/26 R/25 I/324 T/46 H/1 G/1 | - | N:S/5 S:S/3 I:S/1 I:T/6 R:S/2 |
| **NEP** | 67 | - | 0.069 | structure | - | E/859 G/200 K/1 D/10 N/3 | - | E:G/6 E:E/2 D:E/2 D:D/1 D:G/1 G:G/1 |
| **NEP** | 83 | 0.026 | - | overall | - | V/939 M/7 -/19 I/85 L/1 A/1 C/21 | - | M:V/1 V:V/3 I:V/5 I:M/1 C:V/1 C:C/1 |
| **NEP** | 111 | 0.007 | - | - | - | Q/1022 ?/1 L/1 -/26 H/1 P/2 S/19 N/1 | - | P:Q/1 Q:S/1 |
| **NP** | 50 | 0.038 | - | - | S/280 N/1 | S/1904 G/27 N/2 R/68 K/1 -/20 | - | G:S/1 N:S/1 S:S/10 R:S/1 G:R/1 |
| **NP** | 51 | 0.049 | - | - | D/281 | D/1990 N/11 E/1 -/20 | D:D/1 | D:D/4 D:N/2 |
| **NP** | 245 | - | 0.054 | - | S/277 G/4 | S/2021 G/1 | G:S/1 | S:S/2 |
| **NP** | 409 | 0 | - | overall, bipolar, structure, volume, composition, charge | Q/281 | Q/2001 -/15 S/6 | - | Q:Q/4 Q:S/1 |

| | | | | | | | | |
|---|---|---|---|---|---|---|---|---|
| **NP** | 411 | 0.011 | - | overall,composition | T/280 A/1 | T/1993 ?/4 -/23 L/1 A/1 | - | T:T/8 L:T/1 |
| **NP** | 473 | 0.018 | - | - | N/275 S/4 -/2 | N/1963 S/11 -/36 D/9 ?/3 | N:S/1 | N:S/1 D:N/1 |
| **NP** | 495 | 0 | - | overall, volume, composition | E/278 -/3 | E/1895 D/1 -/112 ?/9 R/5 | - | E:E/7 E:R/1 |
| **NS1** | 8 | 0.02 | - | overall, volume, charge | S/144 -/4 | S/1740 -/26 D/2 | - | D:S/1 S:S/4 |
| **NS1** | 20 | 0.012 | - | volume | K/147 -/1 | K/1757 -/5 N/1 G/2 ?/2 E/1 | - | G:K/1 K:K/3 |
| **NS1** | 23 | 0.009 | - | - | A/147 -/1 | A/1736 V/2 -/5 G/1 S/24 | - | A:S/1 S:S/1 |
| **NS1** | 24 | 0.003 | - | structure, composition | D/147 -/1 | D/1737 -/5 N/2 M/24 | - | D:M/1 |
| **NS1** | 27 | 0.035 | - | overall | L/144 M/3 -/1 | L/1678 M/83 Q/2 -/5 | L:M/1 | L:M/9 L:L/4 |
| **NS1** | 48 | 0 | - | overall, bipolar, structure, volume, composition, charge | S/148 | S/1248 I/13 T/1 N/502 -/3 G/1 | - | I:S/2 N:S/8 |
| **NS1** | 82 | 0.001 | - | charge | A/144 -/4 | A/1093 V/3 T/126 G/3 -/525 L/11 S/1 D/5 N/1 | - | A:G/1 A:T/5 A:P/1 L:P/2 A:A/5 A:D/1 |
| **NS1** | 83 | 0 | - | bipolar, structure | P/5 S/138 -/4 F/1 | S/761 P/467 L/2 -/525 K/7 Q/4 A/1 Y/1 | - | P:S/2 S:T/1 K:T/1 K:Q/1 Q:Q/1 S:S/2 |
| **NS1** | 84 | 0.032 | - | - | V/144 -/4 | V/1184 M/18 L/9 -/517 R/2 T/4 ?/6 N/2 K/1 S/25 | - | V:V/5 M:V/2 M:R/1 M:T/2 K:M/1 M:S/1 L:V/2 S:V/1 S:S/1 |
| **NS1** | 86 | 0.033 | - | - | A/136 ?/3 -/4 T/5 | A/1325 T/354 S/5 V/70 I/4 -/6 F/1 ?/1 N/1 D/1 | A:T/2 | A:T/6 A:A/2 A:V/6 I:T/2 T:T/1 A:S/1 |
| **NS1** | 88 | 0.026 | - | - | R/137 H/4 ?/7 | R/1612 H/137 C/11 Y/1 Q/2 ?/5 | H:R/1 | C:R/3 R:R/4 H:R/5 H:Q/1 |
| **NS1** | 91 | 0.013 | - | - | T/138 S/3 ?/7 | T/1754 I/3 ?/5 N/1 S/2 A/3 | S:T/1 | T:T/5 S:T/1 A:T/1 |
| **NS1** | 161 | 0.016 | - | - | S/147 L/1 | S/1696 L/13 T/58 P/1 | - | S:T/3 L:S/1 S:S/1 |
| **NS1** | 166 | 0.022 | - | - | L/148 | L/1579 I/12 F/157 M/19 V/1 | - | I:L/1 F:L/3 L:M/1 I:M/1 |
| **NS1** | 172 | 0.027 | - | - | E/147 ?/1 | E/1732 K/18 A/18 | - | E:K/3 A:E/1 |

| | | | | | | | | |
|---|---|---|---|---|---|---|---|---|
| **NS1** | 176 | 0.025 | - | - | N/147 ?/1 | N/1720 D/34 S/7 I/6 H/1 | - | D:N/1 N:S/1 |
| **NS1** | 189 | 0.02 | - | - | D/145 G/1 N/2 | D/1749 N/6 G/11 Y/1 ?/1 | D:N/1 | D:N/1 D:G/3 D:D/1 |
| **NS1** | 197 | 0 | - | - | T/147 N/1 | T/1615 I/101 A/10 N/39 V/2 ?/1 | - | I:T/6 I:V/1 A:T/2 N:T/1 |
| **NS1** | 202 | 0.01 | - | - | A/145 T/1 -/2 | A/1673 T/90 V/1 S/2 -/2 | - | A:T/5 |
| **NS1** | 205 | 0.032 | - | - | S/146 ?/2 | S/1592 I/24 G/20 ?/3 N/117 D/4 V/7 R/1 | - | I:S/2 G:S/2 S:S/1 N:S/3 D:N/1 D:G/1 I:V/2 |
| **NS1** | 207 | 0.022 | - | - | N/146 ?/2 | N/1085 H/1 ?/3 D/669 G/10 | - | D:N/5 D:G/1 |
| **NS1** | 209 | 0 | - | overall, structure, volume, charge | D/141 G/3 N/2 ?/2 | D/1211 N/129 G/374 S/17 -/1 V/34 ?/2 | D:G/1 | D:N/13 D:G/10 G:S/3 D:V/1 D:D/2 |
| **NS1** | 210 | 0.001 | - | - | G/148 | G/1711 W/3 E/11 R/42 -/1 | - | G:W/1 E:G/2 G:R/3 |
| **NS1** | 212 | 0.018 | - | - | P/147 T/1 | P/1093 L/500 S/7 -/1 F/2 T/163 A/1 H/1 | - | L:P/4 P:T/1 |
| **NS1** | 213 | 0 | - | overall, volume, charge | P/144 S/4 | P/1717 S/33 T/2 L/15 -/1 | P:S/1 | P:S/8 L:P/3 |
| **NS1** | 214 | 0 | - | overall | L/148 | L/1708 F/48 H/10 -/1 Q/1 | - | F:L/8 H:L/1 |
| **NS1** | 215 | 0 | - | - | P/147 S/1 | P/1691 H/2 L/43 S/14 -/1 F/3 T/14 | - | L:P/2 P:S/2 F:L/1 P:T/1 |
| **NS1** | 216 | 0.004 | - | overall, volume | P/147 S/1 | P/1719 Q/2 S/44 -/1 ?/2 | - | P:S/7 |
| **NS1** | 218 | 0 | - | - | Q/148 | Q/1731 -/8 K/3 ?/1 R/1 W/23 P/1 | - | Q:W/2 |
| **NS1** | 219 | 0.012 | - | - | K/148 | K/1647 E/41 -/3 N/77 | - | E:K/3 K:N/2 |
| **NS1** | 226 | 0.001 | - | - | I/147 T/1 | I/1693 ?/1 T/22 V/49 -/2 S/1 | - | I:T/4 I:V/5 |
| **NS1** | 227 | 0.01 | - | - | E/147 G/1 | E/1725 K/8 G/31 -/2 ?/1 V/1 | - | E:K/1 E:G/3 |
| **NS1** | 228 | 0 | - | overall, volume | S/145 P/3 | Y/3 S/1704 P/59 -/2 | P:S/1 | S:Y/1 P:S/9 |
| **PA** | 24 | 0.045 | - | - | Y/56 | Y/2140 H/6 -/16 ?/2 | - | Y:Y/2 H:Y/1 |

| | | | | | | | | |
|---|---|---|---|---|---|---|---|---|
| **PA** | 29 | - | 0.013 | - | K/56 | K/2143 R/3 -/15 G/1 E/1 ?/1 | - | K:K/1 |
| **PA** | 231 | 0.036 | - | - | A/56 | A/2135 T/16 V/9 -/4 | - | A:T/4 A:V/2 |
| **PA** | 261 | 0.049 | - | overall | L/56 | L/2107 M/51 S/2 ?/1 F/1 V/1 Q/1 | - | L:M/4 L:S/1 L:L/7 |
| **PA** | 287 | 0.038 | - | overall | A/56 | A/2138 S/25 G/1 | - | A:A/9 A:S/1 |
| **PA** | 305 | 0.035 | - | - | Y/56 | Y/2148 C/8 H/7 ?/1 | - | C:Y/2 Y:Y/7 H:Y/1 |
| **PA** | 308 | 0.008 | - | - | I/56 | I/2086 S/2 T/47 V/28 L/1 | - | I:S/1 I:T/3 I:I/1 I:V/1 |
| **PA** | 489 | 0.003 | - | overall, bipolar, structure, composition | C/53 S/3 | C/2130 S/13 ?/21 | C:S/1 | C:S/1 C:C/3 |
| **PA** | 544 | 0 | - | overall, bipolar, structure, volume, composition, charge | E/56 | E/2136 K/1 ?/21 Q/1 R/5 | - | E:E/4 E:Q/1 E:R/1 |
| **PA *** | 655 | 0.01 | 0 | - | L/34 F/19 S/3 | L/2141 -/18 ?/1 I/2 F/2 | L:L/1 F:S/1 | L:L/5 F:L/1 |
| **PA** | 714 | 0.044 | - | - | A/56 | A/2123 ?/2 -/27 V/6 T/6 | - | A:V/2 A:T/1 |
| **PB1** | 64 | - | 0.108 | - | P/520 -/1 L/6 S/3 | P/2518 ?/1 -/12 L/7 S/1 H/1 | L:P/1 P:P/3 P:S/1 | P:P/4 |
| **PB1** | 97 | 0.018 | - | overall | E/520 D/10 | E/2498 -/11 A/1 K/19 N/6 D/4 V/1 | E:E/1 D:E/1 | E:E/7 E:N/1 E:K/2 K:N/1 K:K/1 |
| **PB1** | 114 | 0.03 | - | - | V/520 I/10 | V/2480 -/8 I/47 A/2 L/2 D/1 | I:V/1 V:V/1 | V:V/9 I:V/5 L:V/1 |
| **PB1** | 261 | 0.016 | - | composition | S/492 ?/2 G/18 N/1 C/11 T/4 A/2 | S/2343 ?/9 G/104 N/17 C/63 R/3 I/1 | S:T/1 T:T/1 A:T/1 | G:S/2 S:S/4 C:S/2 R:S/1 N:S/1 |
| **PB1** | 378 | 0.006 | - | - | L/528 ?/2 | L/2497 ?/7 M/33 V/1 F/2 | L:L/3 | L:L/16 F:L/1 |
| **PB1** | 578 | 0.028 | - | overall, bipolar, structure, composition, charge | K/516 R/13 -/1 | K/2449 R/69 E/1 -/3 Q/1 N/17 | K:R/2 K:K/1 | K:K/11 K:R/3 R:R/1 K:N/2 |
| **PB1** | 756 | 0 | - | - | Q/520 -/6 R/3 ?/1 | Q/2389 -/83 ?/7 P/60 R/1 | Q:R/1 | Q:Q/5 P:Q/1 P:P/3 |
| **PB2** | 2 | - | 0.075 | - | E/317 -/16 ?/2 | E/2396 ?/2 -/97 G/1 D/1 S/1 T/1 | - | E:E/9 E:T/1 |

| | | | | | | | | |
|---|---|---|---|---|---|---|---|---|
| **PB2** | 4 | 0.001 | - | overall, bipolar, structure, composition, charge | I/321 -/13 T/1 | I/2406 V/2 -/84 ?/3 R/1 L/1 T/2 | - | I:T/1 |
| **PB2 #** | 58 | 0.025 | - | - | T/39 A/295 -/1 | T/2195 A/293 -/7 K/1 S/2 ?/1 | A:T/2 | S:T/1 T:T/4 A:T/4 |
| **PB2** | 64 | 0.022 | - | - | M/334 -/1 | M/2028 ?/2 T/12 -/8 L/4 I/441 V/4 | - | L:M/1 I:M/6 I:L/1 I:T/1 M:V/1 |
| **PB2** | 123 | 0.015 | - | - | E/328 -/7 | E/2453 G/12 -/8 D/24 ?/1 V/1 | - | D:E/2 E:E/2 E:G/2 |
| **PB2** | 127 | 0.041 | - | - | H/328 -/7 | H/2425 Q/3 -/9 N/3 Y/54 ?/3 R/2 | - | H:H/9 H:Y/3 H:Q/1 |
| **PB2** | 220 | 0.002 | - | overall, bipolar, structure | V/328 -/7 | V/2484 G/1 -/8 I/4 K/2 | - | V:V/6 K:V/1 |
| **PB2** | 411 | 0.033 | - | - | I/333 -/2 | I/2413 M/3 L/8 V/74 T/1 | - | I:M/1 I:I/1 I:L/1 I:V/2 V:V/1 |
| **PB2** | 448 | 0.048 | - | - | N/333 ?/2 | N/2480 S/10 H/5 T/2 K/1 ?/1 | - | N:S/3 N:N/6 H:N/1 |
| **PB2** | 463 | 0 | - | - | I/333 V/1 ?/1 | I/2375 V/102 M/18 L/4 | - | I:V/2 I:M/4 |
| **PB2** | 591 | 0.033 | - | - | Q/329 ?/1 R/5 | Q/2494 K/2 -/1 L/2 | Q:R/1 | Q:Q/5 L:Q/1 |
| **PB2** | 607 | 0.021 | - | composition | L/335 | L/2485 -/1 R/1 W/1 V/2 I/9 | - | L:L/9 L:R/1 L:V/1 I:L/1 |
| **PB2** | 615 | 0.039 | - | - | I/335 | I/2470 -/2 V/14 T/4 M/9 | - | I:I/2 I:M/1 I:V/2 I:T/1 |
| **PB2** | 647 | 0.014 | - | - | I/331 -/1 M/1 V/2 | I/2490 ?/1 V/2 M/2 -/2 L/2 | I:V/1 | I:L/1 |
| **PB2 #** | 676 | 0.041 | - | bipolar | T/9 A/313 P/1 -/1 V/10 ?/1 | T/2039 L/2 A/405 -/3 M/27 I/6 Q/1 S/1 D/1 V/11 ?/3 | A:V/1 A:A/1 | T:T/4 L:T/1 A:T/4 I:T/1 I:M/1 |
| **PB2** | 684 | 0.01 | - | - | A/333 -/1 V/1 | A/2423 T/55 V/4 D/1 E/1 -/3 S/10 N/1 ?/1 | - | A:T/6 A:A/3 A:S/2 A:V/1 |
| **PB2** | 759 | 0 | - | overall, bipolar, structure, composition | N/329 ?/1 -/5 | N/2385 -/103 D/2 Y/2 ?/5 I/2 | - | D:N/1 |
